# Unravelling lactate-acetate conversion to butyrate by intestinal *Anaerobutyricum* and *Anaerostipes* species

**DOI:** 10.1101/2020.06.09.139246

**Authors:** Sudarshan A. Shetty, Sjef Boeren, Thi Phuong Nam Bui, Hauke Smidt, Willem M. de Vos

## Abstract

The D-and L-forms of lactate are important fermentation metabolites produced by intestinal bacteria but have been found to negatively affect mucosal barrier function and human health. Of interest, both enantiomers of lactate can be converted with acetate into the presumed beneficial butyrate by a phylogenetically related group of anaerobes, including *Anaerobutyricum* and *Anaerostipes* spp. This is a low energy yielding process with a partially unknown pathway in *Anaerobutyricum* and *Anaerostipes* spp. and hence, we sought to address this via a comparative genomics, proteomics and physiology approach. We focused on *Anaerobutyricum soehngenii* and compared its growth on lactate with that on sucrose and sorbitol. Comparative proteomics revealed a unique active gene cluster that was abundantly expressed when grown on lactate. This active gene cluster, *lctABCDEF*, encodes a lactate dehydrogenase (*lctD*), electron transport proteins A and B (*lctCB*), along with a nickel-dependent racemase (*lctE*) and a lactate permease (*lctF*). Extensive search of available genomes of intestinal bacteria revealed this gene cluster to be highly conserved in only *Anaerobutyricum* and *Anaerostipes* spp. The present study demonstrates that *A. soehngenii* and several related *Anaerobutyricum* and *Anaerostipes* spp. are highly adapted for a lifestyle involving lactate plus acetate utilization in the human intestinal tract.

## Introduction

The major fermentation end products in anaerobic colonic sugar fermentations are the short chain fatty acids (SCFAs) acetate, propionate and butyrate. Whilst all these SCFAs confer health benefits, butyrate is used to fuel colonic enterocytes, and has been shown to inhibit the proliferation and to induce apoptosis of tumour cells (McMillan et al., 2003; Thangaraju et al., 2009) (Topping and Clifton, 2001). Butyrate is also suggested to play a role in gene expression as it is a known inhibitor of histone deacetylases, and recently it was demonstrated that butyrate promotes histone crotonylation *in vitro* (Davie, 2003; Fellows et al., 2018). Related to this ability, exposure to butyrate during differentiation of macrophages was demonstrated to boost antimicrobial activity (Schulthess et al., 2019). In contrast, lactate is also a common end product of anaerobic fermentation in the gut but has no known health benefits. Rather lactate, and notably the D-enantiomer, has been found to be involved in acidosis, reduction of intestinal barrier function in adults and atopic eczema development in children (Ten Bruggencate et al., 2006; Seheult et al., 2017; Wopereis et al., 2018).

Several elegant pioneering studies have shown that butyrate can also be produced by human anaerobes via the conversion of lactate and acetate (Barcenilla et al., 2000; Duncan et al., 2004). This later turned out to be one of major routes for butyrate formation in the gut (Louis and Flint, 2017). However, this relevant metabolic property is only found in a limited number of phylogenetically related species belonging to the genera *Anaerostipes* and *Anaerobutyricum*, the latter being previously known as *Eubacterium hallii* (Shetty et al., 2018). Since the enantiomers D- and L-lactate are end products of fermentation by primary degraders such as *Bifidobacterium* and other (lactic acid) bacteria, these are important cross-feeding metabolites in the diet driven trophic chain existing in the intestinal microbiome (Duncan et al., 2004; Deis and Kearsley, 2012; Belzer et al., 2017). *Anaerostipes caccae* and the two *Anaerobutyricum* species, *A. soehngenii* and *A. hallii* were described to be capable of converting D- and L-lactate plus acetate to butyrate via the acetyl CoA pathway (Barcenilla et al., 2000; Duncan et al., 2004; Louis and Flint, 2017). Growth on lactate poses a major energetic barrier because it is an energy-dependent process with low energy yield since the first step in conversion of lactate to pyruvate in the presence of NAD^+^/NADH is an endergonic reaction (ΔG°’= + 25kJ mol^-1^). Evidence for a molecular mechanism of conversion of lactate to acetate was first demonstrated in the acetogenic model organism *Acetobacterium woodii*, where for every molecules of lactate converted to acetate only 1.5 molecules of ATP were generated (Weghoff et al., 2015). This mechanism, so-called electron confurcation, was suggested to be wide spread in other anaerobic lactate utilizers and hypothesized to be present in intestinal butyrogenic bacteria (Weghoff et al., 2015; Louis and Flint, 2017; Detman et al., 2019). Therefore, using a proteogenomic approach we aimed to investigate whether intestinal bacteria also employ this electron confurcation to convert lactate to butyrate with a focus on *A. soehngenii*. We grew *A. soehngenii* in the presence of three different carbon sources, lactate plus acetate, sucrose and sorbitol. Comparison of proteomic expression data revealed a complete gene cluster, expression of which was induced when *A. soehngenii* was grown in D,L-lactate plus acetate. Investigation of the gene cluster revealed it to be similar to the previously reported gene cluster in *Acetobacterium woodii* involved in the conversion of D,L-lactate to pyruvate, in which an electron transport flavoprotein (EtfAB complex) is active. Extensive search of publicly available bacterial genomes revealed that this gene cluster is highly conserved in several *Anaerobutyricum* and *Anaerostipes* spp. among the butyrate producers from the human intestinal tract. This unique genomic organization suggests that both *Anaerobutyricum* species and *Anaerostipes caccae* and also a recent human infant isolate *Anaerostipes rhamnosivorans* (Bui et al., 2014) have adapted to a lifestyle involving efficient lactate plus acetate utilization in the human intestinal tract.

## Experimental Procedures

### Bacterial strain and growth media

*Anaerobutyricum soehngenii* (DSM17630) strain L2-7 was grown routinely in a medium as described previously (Shetty et al., 2018). The composition of the growth medium was: yeast extract (4.0 g/L), casitone (2.0 g/L), soy peptone (2.0 g/L), NaHCO_3_ (4.0 g/L), KH_2_PO_4_ (0.41 g/L), MgCl_2_.6H_2_O (0.1 g/L), CaCl_2_.2H_2_O (0.11 g/L), cysteine-HCL (0.5 g/L), vitamin K1 (0.2ml), hemin (1ml), and trace elements I, trace elements II and vitamin solutions. The trace elements I (alkaline) solution contained the following (mM): 0.1 Na_2_SeO_3_, 0.1 Na_2_WO_4_, 0.1 Na_2_MoO _4_ and 10 NaOH. The trace elements II (acid) solution was composed of the following (mM): 7.5 FeCl_2_, 1 H_3_BO_4_, 0.5 ZnCl_2_, 0.1 CuCl_2_, 0.5 MnCl_2_, 0.5 CoCl_2_, 0.1 NiCl_2_ and 50 HCl. The vitamin solution had the following composition (g/L): 0.02 biotin, 0.2 niacin, 0.5 pyridoxine, 0.1 riboflavin, 0.2 thiamine, 0.1 cyanocobalamin, 0.1 p-aminobenzoic acid and 0.1 pantothenic acid. This basal medium was supplemented with 30mM of sodium acetate (termed basal medium with acetate). For routine use, the medium was distributed in 35 ml serum bottles sealed with butyl-rubber stoppers and incubated at 37 °C under a gas phase of 1.7 atm (172 kPa) N_2_/CO_2_ (80 : 20, v/v). The pH of the medium was adjusted to 7.0 before autoclaving. The vitamin solution was filter sterilized and added to the media bottles after autoclaving.

### Global proteomic profiling of different growth substrates

Actively growing *A. soehngenii* was pre-cultured in medium with 60 mM glucose and inoculated (5%) in 450 ml of media in triplicates containing 20 mM sucrose, 40 mM D-sorbitol or 80 mM D,L-lactate, respectively. In all three conditions, 30 mM of acetate was added in the medium as it is shown to improve growth of *A. soehngenii* (Duncan et al., 2004; Shetty et al., 2018). Cells were harvested at mid-exponential phase. The samples were centrifuged at 4700 rpm for 30 min at 4°C. The cells and supernatant were stored immediately in -80°C until protein extraction. The supernatant was used to measure SCFAs as well as consumption of sucrose and sorbitol using Shimadzu Prominence-i LC-2030c high-performance liquid chromatography (HPLC). The column used was Shodex SUGAR SH1011. For the mobile phase, 0.01N H_2_SO_4_ was used to which 30 mM of crotonate was added as internal standard.

For whole cell protein extraction, cells were thawed on ice and washed with reduced phosphate buffer solution (PBS with 0.2 mM Titanium (III) citrate) twice before extraction of proteins. The cell pellet was resuspended in SDS-DTT-Tris-Lysis buffer, and cells were disrupted using a French pressure cell (1500 psi). For every sample, the cells were passed through the French pressure cell three times (cell disruption checked under light microscope). After cell disruption, the samples were placed immediately on ice. The extracted proteins were denatured by heat treatment (95°C for 30 min). Protein quantification was done using the Qubit™ Protein Assay Kit (ThermoFisher Scientific, cat.no. Q33211). A total of 50 µg of proteins plus 5 µl of loading dye was mixed and electrophoresed on 10% SDS gels (10% Mini-PROTEAN® TGX™ Precast Protein Gels) for 40 min at 20 mA. Staining was done using Coomassie Brilliant Blue R250 (2.5 g/L) (in 45% methanol and 10% glacial acetic acid) for 3 h and destained with destaining solution (25% methanol and 10% glacial acetic acid in ultrapure water) overnight.

The gels were treated for reduction and alkylation using 50 mM ammonium bicarbonate, 20 mM beta-mercaptoethanol (pH 8) and 20 mM acrylamide (pH 8). Each lane was cut into three even slices, which were further cut into small pieces of approximately 1–2 mm^2^. Trypsin digestion was performed by adding 50 μL of sequencing grade trypsin (5 ng/μL in 50 mM ammonium bicarbonate) and incubated at room temperature overnight while shaking. The resulting tryptic peptide samples were desalted and concentrated using a μColumn (made with LichroprepC18) and subjected to nanoLC-MS/MS using a Proxeon Easy nanoLC and an LTQ-Orbitrap XL instrument (Lu et al., 2011; Wendrich et al., 2017).

### Downstream proteomics data processing, analysis and visualization

Raw data was initially processed using MaxQuant 1.5.2.8 (54) (false discovery rates were set to 0.01 at peptide and protein levels) and additional results filtering (minimally two peptides are necessary for protein identification of which at least one is unique and at least one is unmodified) were performed as described previously (Smaczniak et al., 2012; Cox et al., 2014; Wendrich et al., 2017). Filtered proteinGroups normalized (LFQ) intensities were further analyzed using the DEP bioconductor/R package (Zhang et al., 2018). All proteins that were detected in two out of three replicates of at least one condition were used for further analysis. Missing data was imputed using random draws from a Gaussian distribution centered around a minimal value to control for proteins missing not at random (MNAR). Mostly, these proteins were potentially missed in one condition as a consequence of their intensities being below the detection limit. These pre-processed data were then subjected to variance stabilization, and pair-wise differentially abundant proteins were identified using the limma R package (Ritchie et al., 2015). Here, the tested contrasting conditions were growth on lactate vs sorbitol, lactate vs sucrose, and sorbitol vs sucrose for *A. soehngenii*. Visualization was done using R packages, ggplot2 and ggpubr (Wickham, 2011). The Rmarkdown file with codes to reproduce the analysis of proteomics data obtained from MaxQuant 1.5.2.854 is available at: https://github.com/mibwurrepo/Shetty_et_al_Anaerobutyricum_physiology

### Identification of gene neighbourhood and phylogeny

Gene clusters were identified using the Joint Genome Institute Integrated Microbial Genomes & Microbiome System (JGI-IMG/MER) webserver (Markowitz et al., 2012). The amino acid sequences for EHLA_0974 and EHLA_0978 were searched against the IMG database (Markowitz et al., 2012). The BLASTp hits were limited to only those with an E-value cut-off of 1e-10 and at least 60 % identity. The amino acid sequences were downloaded in fasta format from the IMG website. The sequences were aligned in MEGA6 using ClustalW. The aligned amino acid sequences were used to reconstruct their phylogenetic relationship based on the Jones-Taylor-Thornton (JTT) substitution model.

## Results and Discussion

### Fermentation of carbon sources and global proteomic expression profiles

Actively growing *A. soehngenii* cells pre-cultured on glucose were inoculated in basal medium containing 30mM acetate plus either 20 mM sucrose, 40 mM D-sorbitol or 80 mM D,L-lactate as carbon sources. Acetate was supplied in the media as it has been previously shown to improve growth of *A. soehngenii* (Duncan et al., 2004; Shetty et al., 2018). Acetate was utilized along with other substrates to produce butyrate as the major end product, confirming its importance in supporting growth of *A. soehngenii* (Table 1). High amounts of formate (15.4 ± 0.4 mM, mean ± s.d) were detected from utilization of sucrose (10.9 ± 1.6 mM) while this amount was much lower (5.6 ± 1.7) or none in the case of sorbitol and lactate respectively (Table 1). The ratio for lactate:acetate:butyrate was 2:1:1.5 which is in agreement with previously reported stoichiometry (Duncan et al., 2004). The carbon recovery was 70 %; 64 %; 82.3 % for lactate; sorbitol and sucrose, respectively. The missing carbon can be attributed to biomass production and intermediates that could not be detected by high-performance liquid chromatography at mid-log phase.

**Table 1:**
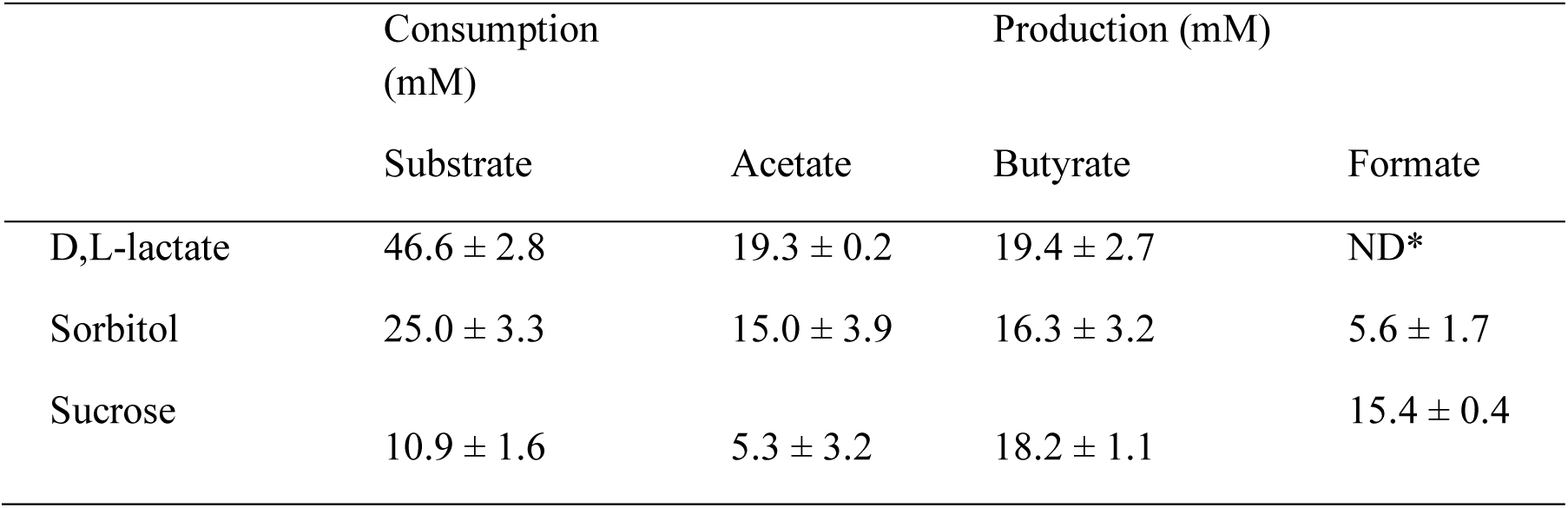
Substrate utilisation and fermentation end products for *A. soehngenii*. **\***ND: not detected. Carbon recovery for D,L-lactate was 70 %, for sorbitol it was 64 % and for sucrose it was 82.3 % at the time of sampling at mid exponential phase. The amount of formed CO_2_ is calculated based on assumption that 1 mole of C6 compound utilized releases 2 moles of CO_2_ or 1 mole formate, hence 1 mole of lactate releases only 1 mole of CO_2_ (Bui et al., 2019)

Next, we compared the total expressed proteome of *Anaerobutyricum soehngenii* when grown on different carbon sources to mid exponential phase. On average, we detected 617 ± 17 proteins present in the minimum of two biological replicates in each condition, of which 453 were detected in all the three growth conditions (Supplementary Figure S1A, B). Principal component analysis of the expressed proteome profiles suggested global differences in protein abundances between the three growth conditions, *i.e.* D,L-lactate, sucrose and sorbitol (Supplementary Figure S1C). Pearson’s correlation between replicates was for all conditions >0.89 (Supplementary Figure S1D).

### Differential proteomes of *A. soehngenii* during growth on sorbitol and sucrose

Sucrose is a common substrate in the distal part of the upper intestinal tract and the ability to utilize sucrose is one of the differentiating features for *A. soehngenii* compared to *A. hallii* (Shetty et al., 2018). Sugar alcohols are commonly used as replacement of glucose for controlling the blood glucose levels in humans. One such common sugar alcohol is sorbitol, which is metabolized slowly by the human body and hence can be available as a carbon and energy source for bacteria in the large intestine (Deis and Kearsley, 2012). Therefore, we sought to identify the active proteins in the metabolic pathway that leads to conversion of sorbitol and sucrose to butyrate and formate.

In total, we identified 23 proteins that were differentially expressed (adjusted *p*-value = 0.05, log_2_-fold change = 1.5) when *A. soehngenii* was grown in presence of sorbitol compared to sucrose. Among these, seven were overexpressed and 16 were less expressed when cells were grown on sorbitol compared to sucrose (Figure 1A). Further inspection of the expressed proteome revealed that *A. soehngenii* has an inducible operon for sorbitol uptake. The genes encoding the phosphotransferase system (PTS) for sorbitol are located between the *gutM* gene coding for the glucitol/sorbitol operon activator and *licR* encoding the lichenan operon transcriptional antiterminator (Figure 1B). The significantly higher (log_2_ fold change, log_2_FC > 6) abundance of these proteins during growth on sorbitol when compared to that on sucrose suggests that this gene cluster is coordinately expressed and highly induced in the presence of sorbitol. The promoter region is located before the gene EHLA_1883 that encodes sorbitol-6-phosphate dehydrogenase, while the terminator is located immediately after the transcriptional terminator EHLA_1878.

**Figure 1:**
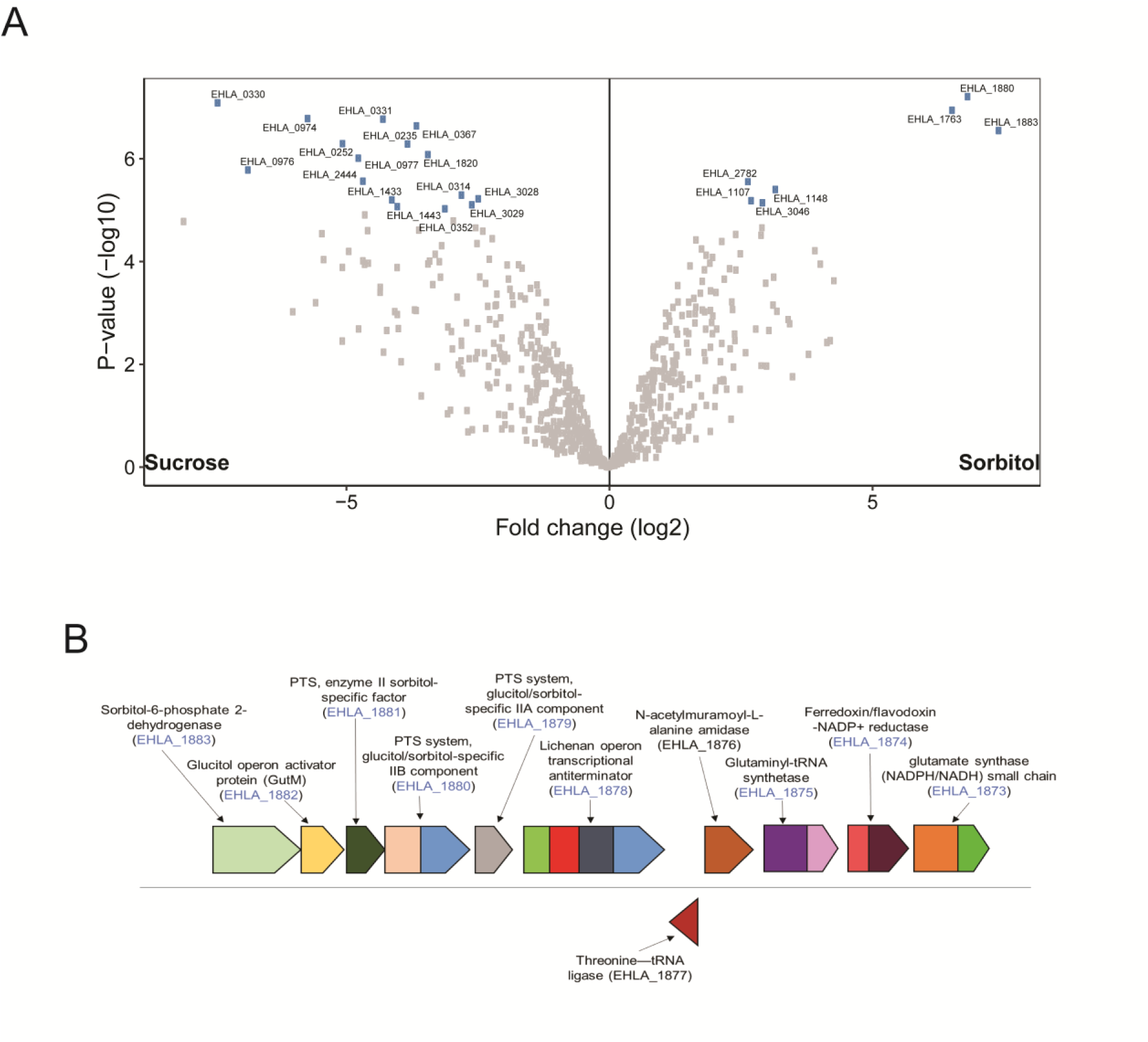
Comparison of *A. soehngenii* expressed proteomes when grown on sorbitol or sucrose. **A]** Volcano plot depicting the differential protein expression of *A. soehngenii* when grown on sucrose or sorbitol. The significantly different (adjusted *p*-value < 0.05) proteins are labelled with corresponding locus tags. **B]** Organisation of gene cluster involved in sorbitol utilization in the genome of *A. soehngenii*. The locus tags detected in the proteome are coloured in blue.

Comparison of the expressed proteome of cells grown on sucrose and sorbitol further revealed that the product of EHLA_0352, bacterial extracellular solute-binding protein (IMG-annotation: carbohydrate ABC transporter substrate-binding protein, CUT1 family (TC 3.A.1.1.-)) had a ten-fold higher abundance in cells grown on sucrose (Supplementary table S1). We identified two locus tags EHLA_0193 and EHLA_0331 encoding a protein-Npi-phosphohistidine-sugar phosphotransferase (PTS) system. The protein encoded by EHLA_0331 was less abundant (log_2_FC >4, adjusted *p*-value = 0.039) in sorbitol compared to sucrose. Also the protein encoded by EHLA_0193 was observed at lower abundance (log_2_FC >8, adjusted p-value = 0.074), albeit not significantly, but we observed that it was >5 log2 fold higher in abundance in cells grown on sucrose compared to cells grown either on D,L-lactate or sorbitol (Supplementary table S1). Upstream of the gene encoding the sucrose PTS is a LacI-type HTH domain containing gene (EHLA_0192), the product of which was not detected in the proteome data but is likely a transcriptional regulator controlling the expression of the sucrose-PTS gene. Next, we did a BLASTp search for the active proteins involved in sucrose utilization in the publicly available genome of *A. hallii* DSM3353. The products encoded by the *A. soehngenii* genes with locus tags EHLA_0192 and EHLA_0193 were not identified in *A. hallii*, while a homologue of the protein encoded by the gene with locus tag EHLA_0195 was identified, which shared 52% identity with a 4-alpha-glucanotransferase in *A. hallii.* Additionally, the gene encoding maltose 6’-phosphate phosphatase (EHLA_0194) did not have a homologue in the genome of *A. hallii*. Based on our findings, we hypothesize that genes with the locus tags EHLA_0192, EHLA_0193, EHLA_0194, and EHLA_0195 are likely responsible for the ability of *A. soehngenii* to utilize sucrose. These genes are not (completely) present in *A. hallii*, which cannot grow on sucrose (Shetty et al., 2018).

### Proteomics-guided identification of the active D,L-lactate utilization gene cluster

Several intestinal bacteria such as those belonging to the genera *Bifidobacterium* and *Lactobacillus* are known lactate producers. Lactate is one of the major cross-feeding metabolites that is converted to butyrate by butyrate producing bacteria such as *A. soehngenii* (Duncan et al., 2004). All known bacteria belonging to the genera *Anaerobutytricum* and *Anaerostipes* have been shown to convert lactate and acetate into butyrate but only *A. soehngenii* and *A. hallii* as well as *Anaerostipes caccae* and *Anaerostipes rhamnosivorans* have been described to utilize both D and L-forms of lactate (Duncan et al., 2004). However, to the best of our knowledge, the active genes involved in butyrogenesis from D,L-lactate and acetate have not been investigated in detail for members of *Anaerobutyricum* and related genera. The expressed proteome of *A. soehngenii* revealed that 31 and 39 proteins were significantly increased when grown on D,L-lactate vs sorbitol and D,L-lactate vs sucrose, respectively (log_2_FC >= 1.5, adjusted *p*-value = 0.05). A total of 15 proteins were shared between the two comparisons (including those encoded by genes with locus tags EHLA_0973, EHLA_0974, EHLA_0976, EHLA_0977, EHLA_0978, EHLA_0979) that were induced by growth on D,L-lactate. Proteins with significantly higher abundance during growth on D,L-lactate included lactate permease, lactate dehydrogenase, electron transfer flavoprotein beta subunit, electron transfer flavoprotein alpha subunit, short-chain acyl-CoA dehydrogenase, and lactate racemase (encoded by genes with locus tags EHLA_0973 - EHLA_0978) (Figure 2A and B). Remarkably, lactate dehydrogenase (encoded by the gene with locus tag EHLA_0974) co-occurs with EtfA and EtfB, which suggests the presence of a D-lactate dehydrogenase/electron-transferring flavoprotein (LDH/Etf) complex allowing growth on lactate, which is a low energy substrate (Weghoff et al., 2015). Interestingly, a gene coding for a short-chain acyl-CoA dehydrogenase, orthologous to that coding for butyryl–CoA dehydrogenase, was identified to be located downstream of the gene encoding lactate dehydrogenase, and corresponding gene products were found produced together with other lactate-specific proteins. The organisation of the unique D,L-lactate utilization gene cluster, *lctABCDEFG* is depicted in Figure 3A.

**Figure 2:**
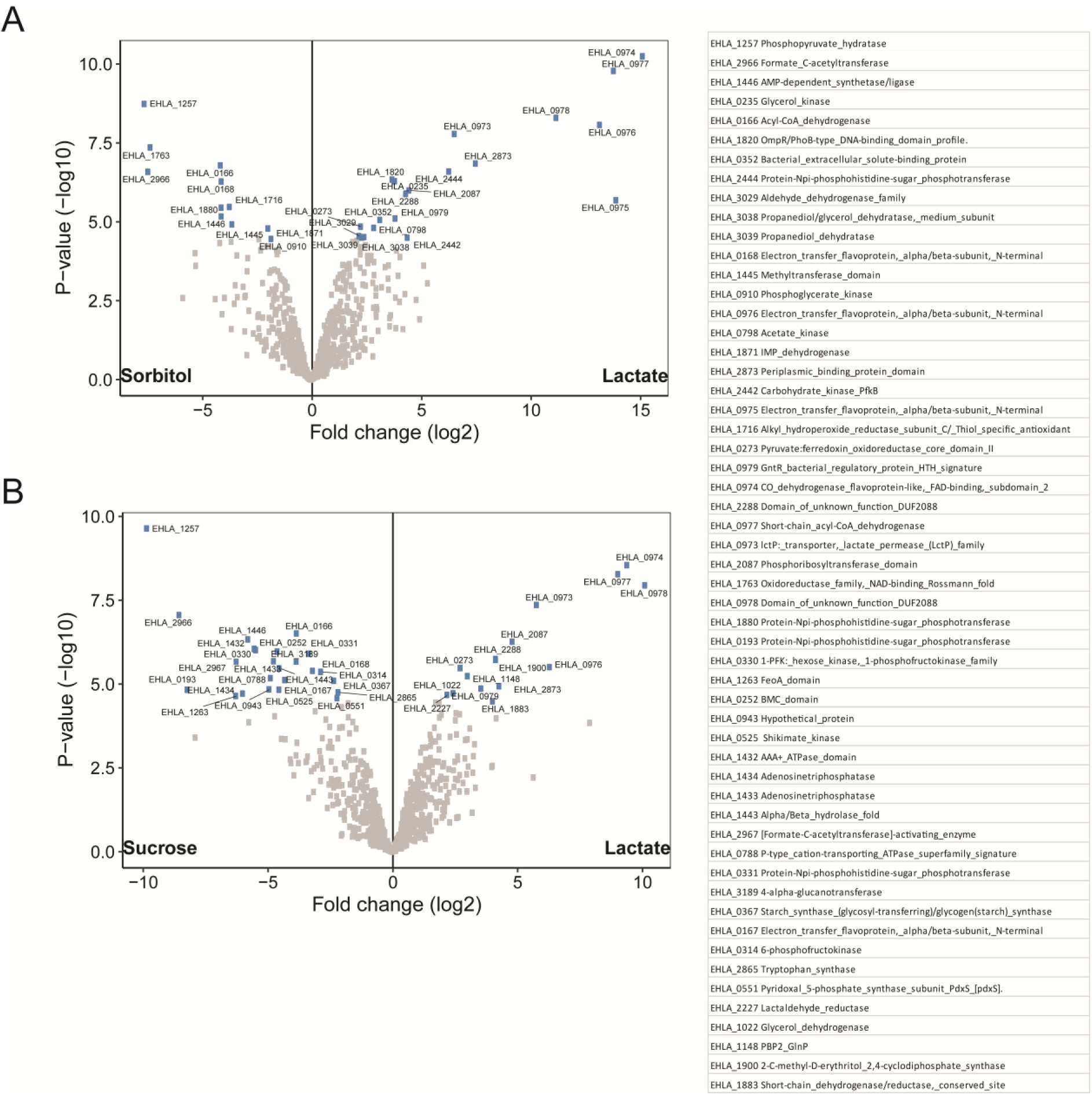
Comparison of *A. soehngenii* expressed proteomes when grown on D,L-lactate compared to sorbitol or sucrose. The significantly different (adjusted *p*-value < 0.05) proteins are labelled with corresponding locus tags. **A]** Volcano plot depicting the differential protein expression of *A. soehngenii* when grown on sorbitol or D,L-lactate. B] Volcano plot depicting the differential protein expression of *A. soehngenii* when grown on sucrose or D,L-lactate. The proteins encoded by genes with the locus tags are indicated.

**Figure 3:**
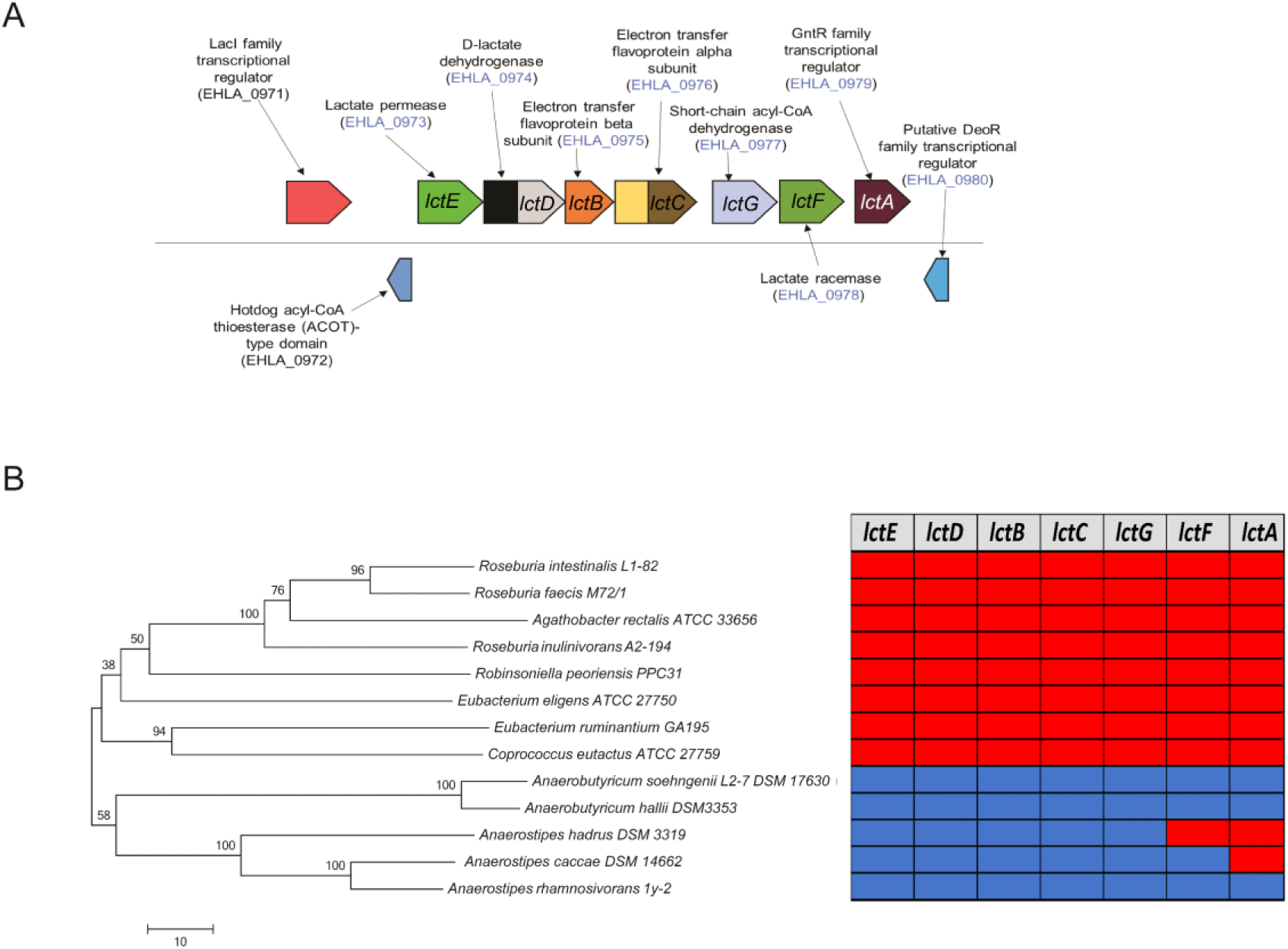
D,L-lactate utilization gene cluster. **A]** Organisation of gene cluster involved in D,L-lactate utilization in the genome of *A. soehngenii*. The locus tags detected in the proteome are coloured in blue. **B]** Phylogenetic tree based on the 16S rRNA gene sequences for *A. soehngenii* and closely related species. The evolutionary history was inferred using the Neighbour-Joining method based on 1000 bootstrap replicates. The amino acid sequences for D-lactate dehydrogenase (EHLA_0974), lactate permease (EHLA_0973) and nickel dependent lactate racemase (EHLA_0978) were searched against the genomes of closely related species. Presence in genome is indicated by blue coloured cells, absence by red coloured cells.

### Comparison of genomes of *Anaerobutyricum* and related species

Next, we searched for homologues of genes encoding lactate permease, lactate dehydrogenase and lactate racemase (encoded by genes with the locus tags EHLA_0973, EHLA_0974 and EHLA_0978, respectively) in the genomes of bacteria closely related to *Anaerobutyricum* (Figure 3B). *A. hallii, Anaerostipes caccae* and *Anaerostipes rhamnosivorans* have homologues of genes coding for lactate dehydrogenase, lactate racemase and lactate permease. *A. rhamnosivorans*, which was previously isolated from infant feces and known to have the ability to utilize D,L-lactate and acetate, has all the genes of the *lct* operon as those found in *Anaerobutyricum* species (Bui et al., 2014). However, the genome of *Anaerostipes hadrus* lacked the gene encoding lactate racemase, which is consistent with its inability to utilize L-lactate (Figure 3B) (Duncan et al., 2004; Allen-Vercoe et al., 2012). We further expanded our search to other bacterial genomes present in the IMG database (Markowitz et al., 2012). The lactate racemase encoded in the genome of *Anaerobutyricum* is predicted to be a nickel-dependent enzyme (Desguin et al., 2014). Further support for the nickel dependency derived from the presence of a set of auxiliary enzymes namely, *larE* (pyridinium-3,5-biscarboxylic acid mononucleotide sulfurtransferase [EC:4.4.1.37], EHLA_1081), *larC1* (pyridinium-3,5-bisthiocarboxylic acid mononucleotide nickel chelatase [EC:4.99.1.12], EHLA_1342) and *larC2* pyridinium-3,5-biscarboxylic acid mononucleotide synthase [EC:2.5.1.143], EHLA_1341), predicted to be involved in nickel incorporation were detected in the proteome but encoded in a scattered way in the genome. A BLASTp search for amino acid sequences similar to the product of the gene with locus tag EHLA_0978 revealed their presence in genomes of members of genera *Intestinimonas* and *Flavonifractor*, as well as species *Eubacterium barkeri, E. aggregans, Pseiudoramibacter alactolyticus, Clostridium merdae, Clostridium jeddahense* and related species. A phylogenetic analysis suggests that lactate racemase (encoded by the gene with locus tag EHLA_0978) in *A. soehngenii* shares similarity to the lactate racemases present in related *E. barkeri, E. aggregans, Intestinimonas* spp. and *Flavonifractor* (Figure 4). *E. aggregans* (isolated from methanogenic bioreactors) and *Intestinimonas* spp. are known for their ability to convert lactate plus acetate to butyrate (Tahar Mechichi and Woo, 1998; Kläring et al., 2013) (Bui et al 2016), although the growth of the human gut isolate *I. butyriciciproducens* AF211 is relatively slow on lactate and acetate (Bui et al., 2015). *Flavonifractor* species are closely related to *Intestinimonas* and can potentially utilize lactate and acetate to produce butyrate.

**Figure 4:**
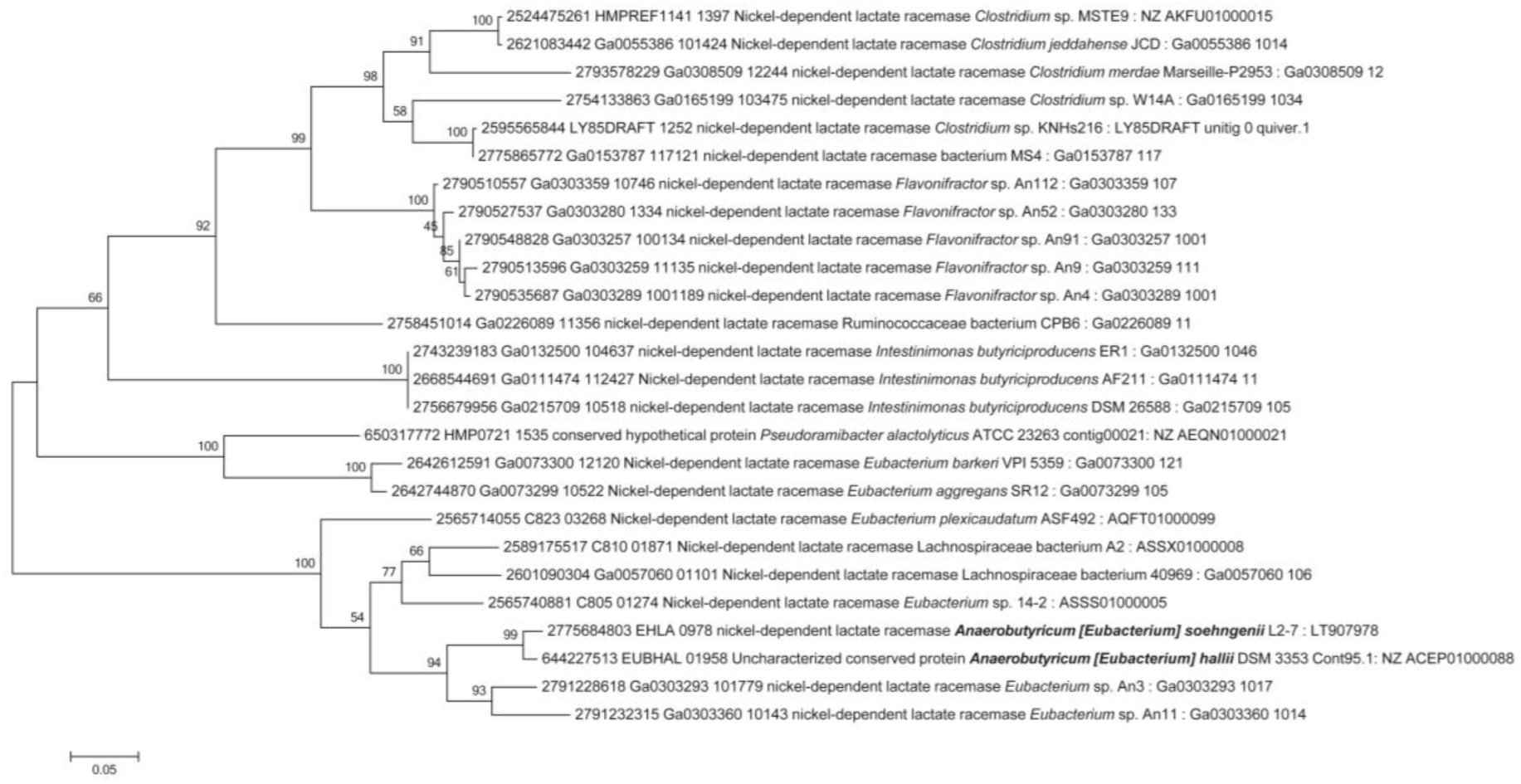
Maximum likelihood phylogenetic tree of 26 lactate racemase homologues. Phylogenetic tree of 26 lactate racemase homologues identified by searching against 55,499 isolate genomes of the IMG/ER database (as of 2 October 2018). Labels represent the IMG gene id, locus tag, IMG annotation for the gene product, taxonomic identity, strain name, and assembly and/or contig/scaffold and NCBI/DDJB/ENA accession number of genomes which contains this gene. The numbers on branches represent the bootstrap values from 1000 replicates. See methods for details on calculation.

A BLASTp search for amino acid sequences of lactate dehydrogenase (encoded by the gene with locus tag EHLA_0974) revealed a wide-spread distribution of homologues in the IMG genome database (Supplementary Figure S2). As with lactate racemase, the phylogenetically related (based on 16S rRNA gene) *Anaerobutyricum* and *Anaerostipes* did not share high identity with each other with respect to lactate dehydrogenase encoding genes and were phylogenetically placed in two distinct clusters. However, further analysis of neighborhood genes indicated that both the *Anaerobutyricum* and *Anaerostipes* lactate dehydrogenase gene is followed by genes coding for electron transfer proteins (Supplementary Figure S3). Overall, these observations provide evidence for unique adaptation of *A. soehngenii, A. hallii* and *Anaerostipes caccae* to utilize an important but low energy cross-feeding metabolite, D,L-lactate.

## Concluding remarks

The genomic and proteomic data presented in this study allowed elucidating key metabolic features of *A. soehngenii* (Figure 5). We identified an inducible operon encoding a sorbitol transporter, which was abundant in the expressed proteome of *A. soehngenii* when grown in presence of sorbitol. The ability to utilize sucrose is likely conferred by two proteins that are not encoded in the genome of the phylogenetically related *A. hallii*, viz. a sucrose PTS and LacI-type HTH domain, which both were highly induced in presence of sucrose. The specialized ability of *A. soehngenii* to convert D,L-lactate and acetate to butyrate is conferred by the unique genomic organization of a lactate utilization gene cluster. Although initial steps of glycolysis require ATP, the overall conversion of glucose to pyruvate is exergonic (Table 2, equation 1). However, growth on lactate poses a major energetic barrier because lactate has to be converted to pyruvate in an NADH-generating and endergonic process (Table 2, equation 2a). The free energy resulting from the coupling with oxidation of ferredoxin has been calculated as -9.5 kJ/mol (see equation 2b (Weghoff et al., 2015)).

**Table 2:** Calculated Gibbs’ free energy for conversion of glucose to pyruvate and lactate to pyruvate.

| Reaction | $\Delta G^0$ |
| --- | --- |
| <u>Equation 1:</u> | -88 kJ/mol |
| Glucose + 2Pi + 2ADP + 2NAD<sup>+</sup><br>→ 2Pyruvate + 2ATP + 2NADH + 2H<sup>+</sup> + 2H<sub>2</sub>O |  |
| <u>Equation 2a:</u> | +25 kJ/mol |
| Lactate + NAD<sup>+</sup> ↔ Pyruvate + NADH |  |
| <u>Equation 2b as proposed by (Weghoff et al., 2015):</u> | -9.5 kJ/mol |
| Lactate + Fd<sup>2-</sup> + 2NAD<sup>+</sup> → Pyruvate + Fd + 2NADH |  |
| <u>Equation 3</u> | -139 kJ/mol |
| 4 Lactate + 2 Acetate → 3 Butyrate + 4 CO<sub>2</sub> + 2H<sub>2</sub> + 2H<sub>2</sub>O |  |
| <u>Equation 4</u> | -249.96 kJ/mol |
| Glucose + 2H<sub>2</sub>O → Butyrate + 2 CO<sub>2</sub> + 4H<sub>2</sub> |  |

**Figure 5:**
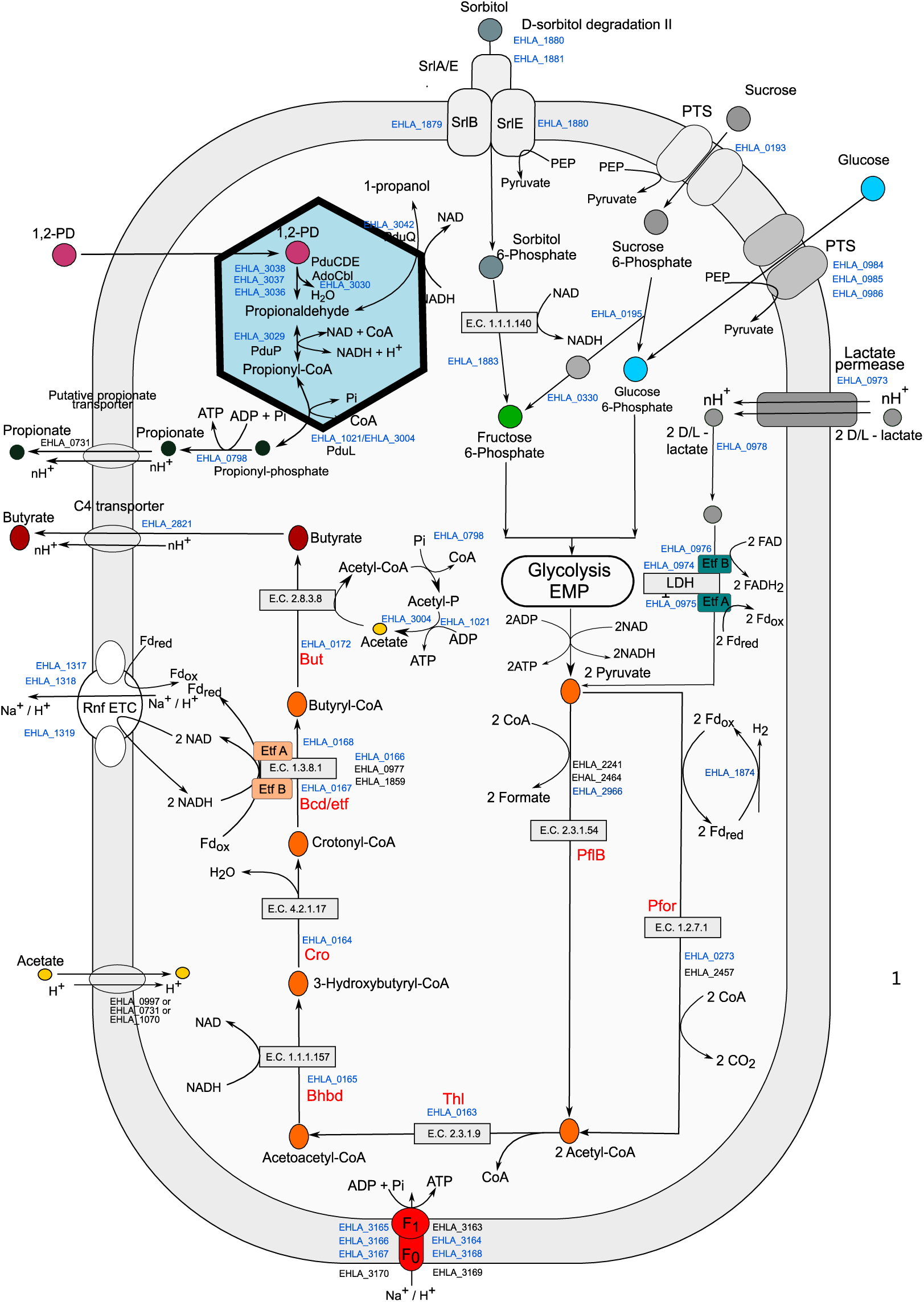
Overview of the key metabolic routes of butyrate and propionate formation in *A. soehngenii*. Relevant locus tags with the prefix EHLA_ are indicated and those in blue relate to gene products that are detected in proteome, while those in black are only identified in genome but not detected in the proteome at any of the growth conditions. The SCFA transporters most likely belong to the C4 TRAP family of transporters. The symport is with a proton and n indicates the number of protons transported across the cell membrane which may vary under various growth conditions.

The bacterial species, *Anaerostipes caccae, A. rhamnosivorans* and the two *Anaerobutyricum* species, *A. soehngenii* and *A. hallii*, are capable of converting D,L-lactate plus acetate to butyrate. Other common intestinal bacteria such as *Faecalibacterium prausnitzii, Roseburia intestinalis* and *Agathobacter rectalis* are known to consume acetate for the production of butyrate from sugars (Duncan et al., 2002; Duncan and Flint, 2008; Heinken et al., 2014). As a consequence, a community with co-occurrence of acetate utilizers will have high competition for acetate, which poses another challenge for the efficient utilization of lactate (along with the low energy yield) by *Anaerobutyricum* and *Anaerostipes* spp. Genome analysis demonstrated that *Anaerobutyricum hallii, A. soehngenii, Anaerostipes caccae* and *Anaerostipes rhamnosivorans* have well-adapted lactate utilization gene clusters that encode all necessary proteins (transcriptional regulator, transporter, racemase (two copies), LDH/Etf complex, a homolog of butyryl-CoA dehydrogenase) that were also detected in high amounts in the expressed proteome data. In conclusion, data presented here has revealed how *Anaerobutyricum soehngenii* and related butyrate producers can overcome the energetic barrier to utilize D,L-lactate to produce butyrate.

## Acknowledgement

We thank Dr. Irene Sanchez Andrea for useful discussions on anaerobic metabolism and thermodynamics during the course of the study and Ton van Gelder for technical support. This research was partly supported by the Netherlands Organization for Scientific Research, Spinoza Award and SIAM Gravity Grant 024.002.002 to WMdV and the UNLOCK project NRGWI.obrug.2018.005 to HS.

