## Supplementary Figure for "Unravelling lactate-acetate conversion to butyrate by intestinal *Anaerobutyricum* and *Anaerostipes* species"

**Competing interests:** The authors declare no competing interests.

**Running title:** Lactate utilization by *Anaerobutyricum* and related species

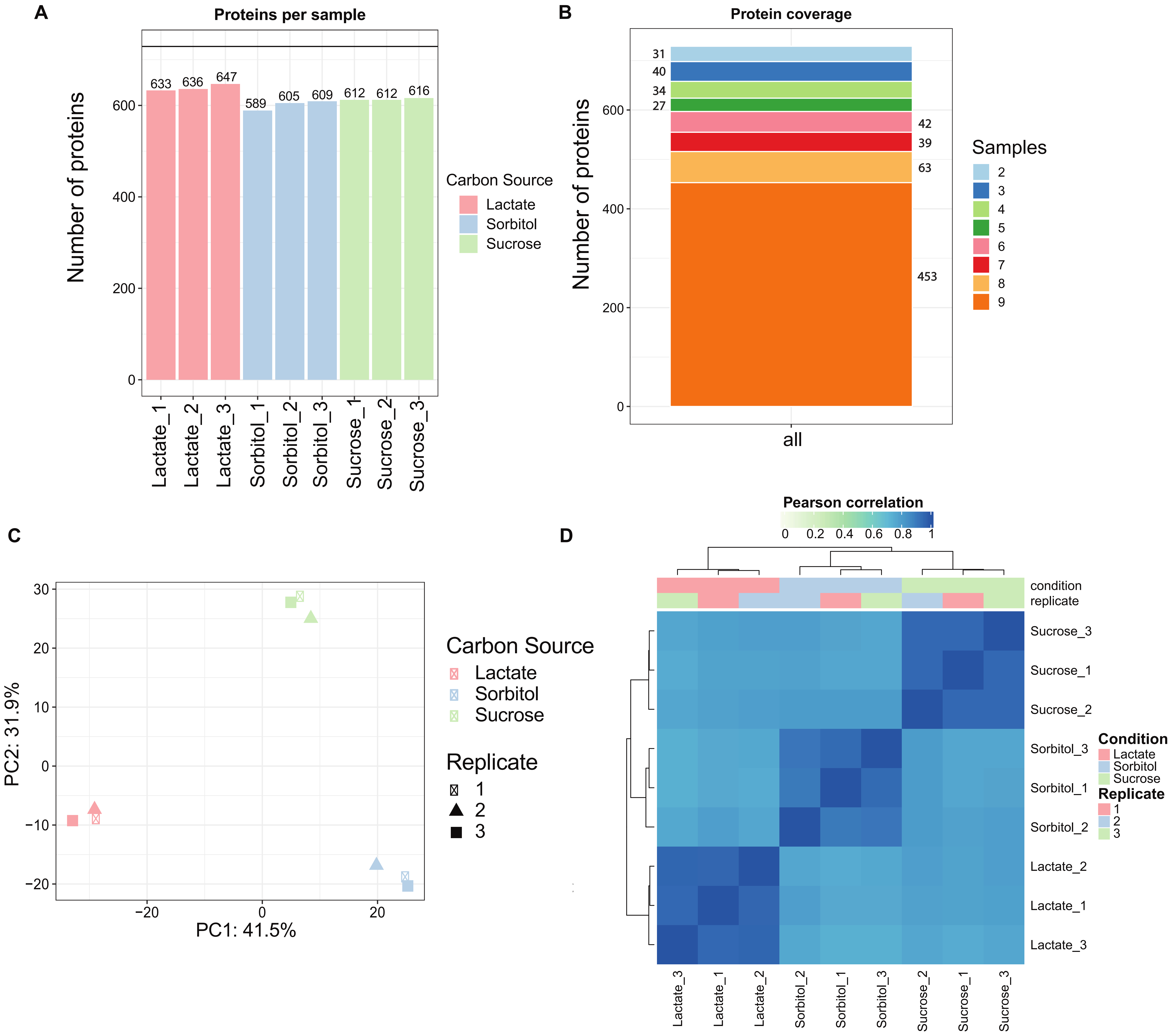


**Figure S1: Overview of the global proteomic comparison.**

A] Number of proteins detected in each of the biological triplicates for different carbon sources. B] Coverage of proteins detected in all the samples. Total nine samples were processed for proteomics, for each carbon source there were triplicates (i.e. 3×3), and the numbers given on either side of the bar plot represent the coverage of proteins in different number of samples. C] Principal components analysis of demonstrating the variation in protein expression profiles under different growth conditions. D] Correlation heatmap depicting the replicability between biological triplicates for each growth condition.

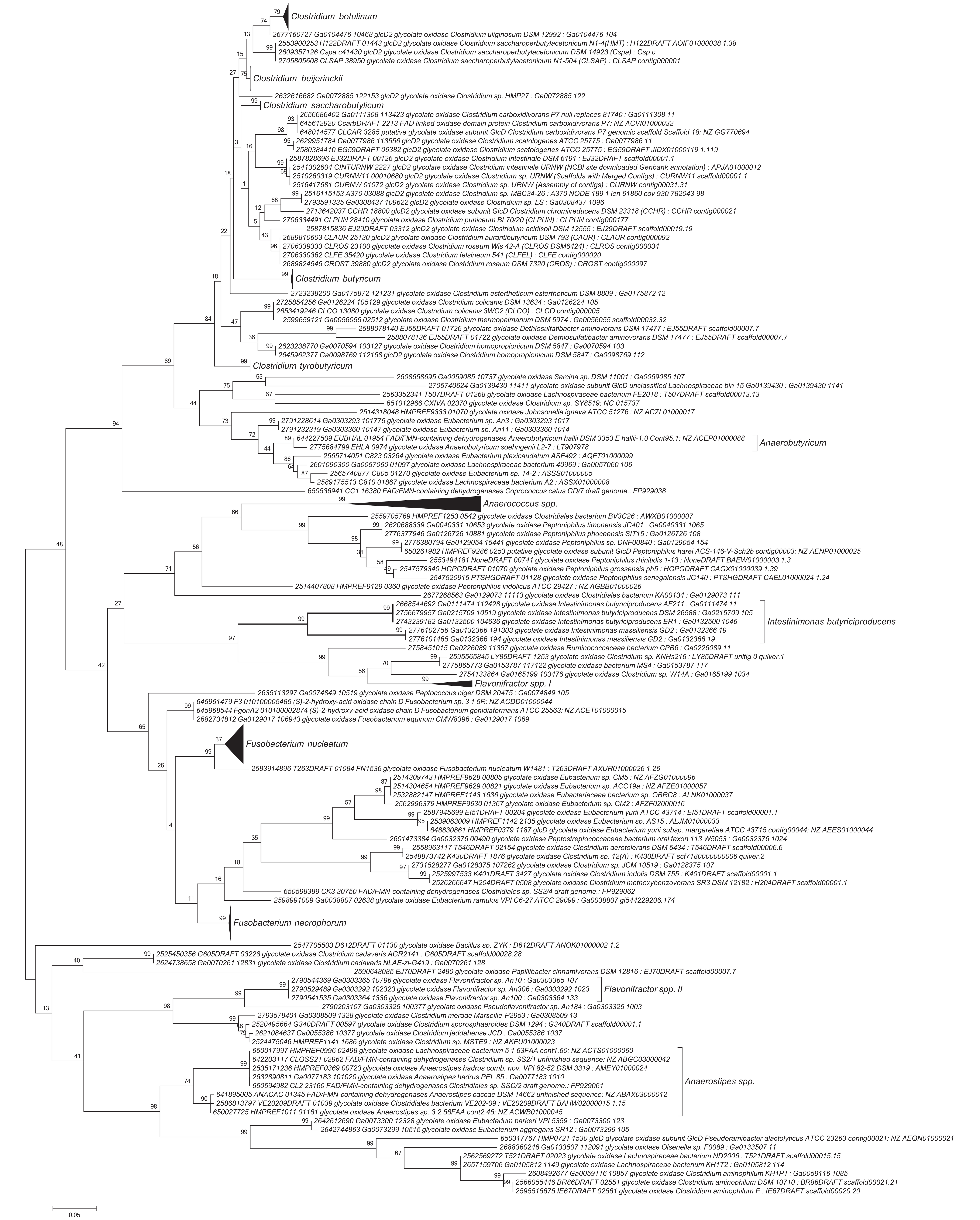


**Figure S2:** Maximum likelihood phylogenetic tree of 285 lactate dehydrogenase homologues. Phylogenetic tree of 285 LDH homologs identifed by searching against 55,499 isolate genomes of the IMG/ER database (as of 2 October 2018). Labels represent the IMG gene id, IMG annotation for the gene product, taxonomic identity, strain name, an assembly and/or contig/scafold which contains this gene. Where several genomes for a species were present, the branch of the tree was collapsed for clarity. The numbers on branches represent the bootstrap values from 1000 replicates. See methods for details on calculation.

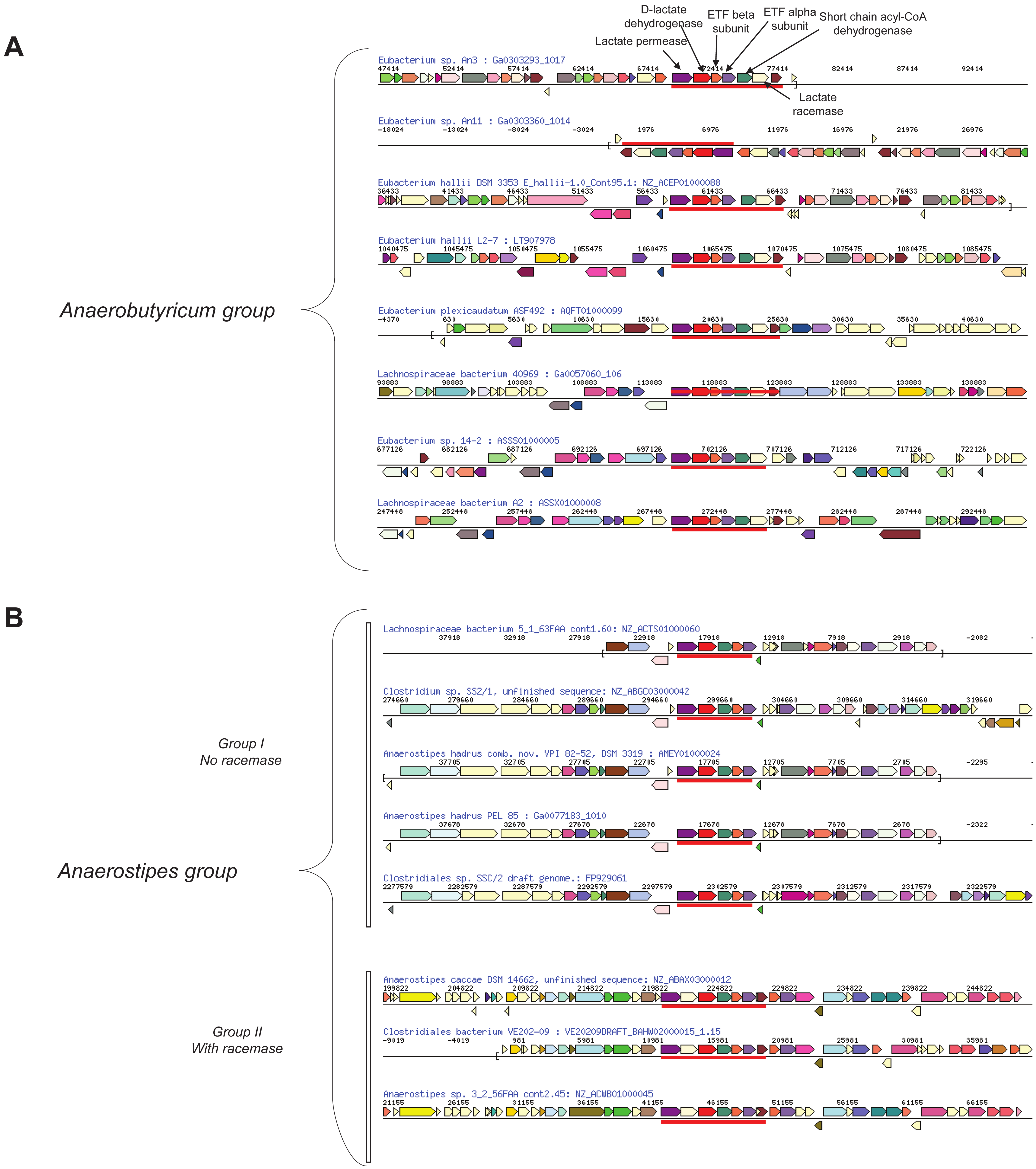


**Figure S3: Comparison of the neighbourhood of genes involved in lactate utilization in genomes of *A. soehngenii* and related lactate utilizing bacteria.** The red line highlights the gene cluster involved in lactate utilization. Neighbourhoods of genes in other genomes with the same top cluster of orthologs (COG) hit and roughly same matching length are shown below using the IMG gene neighbourhood search. Genes of the same colour (except light yellow) are from the same orthologous group (top COG hit). A] Shows selected bacterial genomes that share similar gene organisation as A. soehngenii lactate utilization gene cluster. B] Shows selected bacterial genomes that share similar gene organisation as Anaerostipes caccae lactate utilization gene cluster.
